# annoFuse: an R Package to annotate, prioritize, and interactively explore putative oncogenic RNA fusions

**DOI:** 10.1101/839738

**Authors:** Krutika S. Gaonkar, Federico Marini, Komal S. Rathi, Payal Jain, Yuankun Zhu, Nicholas A. Chimicles, Miguel A. Brown, Ammar S. Naqvi, Bo Zhang, Phillip B. Storm, John M. Maris, Pichai Raman, Adam C. Resnick, Konstantin Strauch, Jaclyn N. Taroni, Jo Lynne Rokita

## Abstract

**Background:** Gene fusion events are a significant source of somatic variation across adult and pediatric cancers and are some of the most clinically-effective therapeutic targets, yet low consensus of RNA-Seq fusion prediction algorithms makes therapeutic prioritization difficult. In addition, events such as polymerase read-throughs, mis-mapping due to gene homology, and fusions occurring in healthy normal tissue require informed filtering, making it difficult for researchers and clinicians to rapidly discern gene fusions that might be true underlying oncogenic drivers of a tumor and in some cases, appropriate targets for therapy.

**Results:** We developed *annoFuse*, an R package, and *shinyFuse*, a companion web application, to annotate, prioritize, and explore biologically-relevant expressed gene fusions, downstream of fusion calling. We validated *annoFuse* using a random cohort of TCGA RNA-Seq samples (N = 160) and achieved a 96% sensitivity for retention of high-confidence fusions (N = 603). *annoFuse* uses FusionAnnotator annotations to filter non-oncogenic and/or artifactual fusions. Then, fusions are prioritized if previously reported in TCGA and/or fusions containing gene partners that are known oncogenes, tumor suppressor genes, COSMIC genes, and/or transcription factors. We applied *annoFuse* to fusion calls from pediatric brain tumor RNA-Seq samples (N = 1,028) provided as part of the Open Pediatric Brain Tumor Atlas (OpenPBTA) Project to determine recurrent fusions and recurrently-fused genes within different brain tumor histologies. *annoFuse* annotates protein domains using the PFAM database, assesses reciprocality, and annotates gene partners for kinase domain retention. As a standard function, *reportFuse* enables generation of a reproducible R Markdown report to summarize filtered fusions, visualize breakpoints and protein domains by transcript, and plot recurrent fusions within cohorts. Finally, we created *shinyFuse* for algorithm-agnostic interactive exploration and plotting of gene fusions.

**Conclusions:** *annoFuse* provides standardized filtering and annotation for gene fusion calls from STARFusion and Arriba by merging, filtering, and prioritizing putative oncogenic fusions across large cancer datasets, as demonstrated here with data from the OpenPBTA project. We are expanding the package to be widely-applicable to other fusion algorithms and expect *annoFuse* to provide researchers a method for rapidly evaluating, prioritizing, and translating fusion findings in patient tumors.

## Background

Gene fusions arise in cancer as a result of aberrant chromosomal rearrangements or defective splicing, bringing together two unrelated genes that are then expressed as a novel fusion transcript (1). Detection of therapeutically-targetable fusion calls is of clinical importance and computational methods are constantly being developed to detect these events in real-time. Recent comparative studies show low concordance of fusion predictions across methods (2), suggesting that many predictions may not represent true events. Additionally, transcriptional read-throughs (3), in which the polymerase machinery skips a stop codon and reads through a neighbouring gene, as well as fusions that involve non-canonical transcripts or gene-homologs, are prevalent in disease datasets, yet the biological relevance of such events is still unclear. This makes it difficult for both researchers and clinicians to prioritize disease-relevant fusions and discern the underlying biological mechanisms and thus, appropriate fusion-directed therapy. Gene fusion events leading to gain-of-function or loss-of-function in kinases and putative tumor suppressor genes, respectively, have been shown to be oncogenic drivers with therapeutic potential, especially in pediatric tumors (4–6). For example, the recurrent fusion *KIAA1549-BRAF* is found across 66-80% of low grade gliomas and results in a fusion transcript that has constitutive BRAF kinase activity (7). *EWSR1-FLI1* is found in nearly 100% of Ewing’s sarcoma and forms an oncogenic RNA complex, driving tumorigenesis (8). To capture highly recurrent and validated fusions such as these, the fusion databases ChimerDB (9) and TumorFusions (10) were developed from RNA fusions called in The Cancer Genome Atlas (TCGA) (11,12) samples. In such large-scale cancer studies, a single algorithm was routinely used to detect fusion calls because using multiple callers often adds complexity of annotation and integration. However, it is now common practice to incorporate data from multiple algorithms to reliably define the fusion landscape of cancers. Recent efforts have reported the importance of using systematic filtering and aggregation of multiple fusion callers to expand the number of biologically-relevant fusions in adult cancers (12,13). However, to our knowledge there are no tools or packages developed to filter, aggregate, and detect recurrent and putative oncogenic fusions in a systematic, flexible, and reproducible manner. Despite the existence of a few tools with working open-source code which can assist in fusion annotation or prioritization, only three are algorithm-agnostic with the remaining tools relying on outdated fusion algorithms, rendering them unusable on current gold standard tools to date, such as STAR-Fusion (14) and Arriba (15) (**Table 1**).

Here, we developed *annoFuse* for annotation, prioritization, and exploration of putative oncogenic gene fusions. We performed technical validation using two independent RNA-Sequencing datasets (TCGA and Pediatric Preclinical Testing Consortium), and finally, applied *annoFuse* to gene fusion calls from STAR-Fusion and Arriba for 1,028 pediatric brain tumor samples provided as part of the OpenPBTA Project (16). To achieve this, we used FusionAnnotator on raw fusion calls to identify and filter red flag fusions, that is, fusions found in healthy tissues or in gene homology databases. Using *annoFuse*, we remove fusions known or predicted to be artifactual and retain high-quality fusion calls. Second, fusions that pass quality checks are annotated (**Table S1**) as previously found within TCGA (10) and each gene partner is annotated as an oncogene, tumor suppressor (17,18), kinase (19), transcription factor (20), and whether it has been reported in the Catalogue of Somatic Mutations in Cancer (COSMIC) Cancer Gene Census (21). In addition, we added the following genes specific to pediatric cancer from literature review: *MYBL1* (22), *SNCAIP* (23), *FOXR2* (24), *TTYH1* (25), and *TERT (26–29)* as oncogenes and *BCOR* (30) and *QKI* (4) as tumor suppressors. Finally, we determined the recurrence pattern for fusions across the cohort and also recurrently-fused genes within each cancer histology.

## Implementation

We implemented *annoFuse* using the R programming language R version 4.0.2 (2020-08-13). The R packages required to install and run *annoFuse* are reshape2, dplyr, tidyr, ggplot2, qdapRegex, ggpubr, tibble, ggthemes, EnsDb.Hsapiens.v86, grid, readr, grDevices, stats, utils, stringr, shiny, shinydashboard, rintrojs, shinythemes, DT, rmarkdown, and methods, with the optional package: knitr. We also created an interactive web-based application of *annoFuse* called *shinyFuse* using the R/Shiny framework.

### R package overview

The *annoFuse* package was developed to provide a standardized filtering and annotation method for fusion calls from Arriba and STAR-Fusion, first and second place winners of the 2017 DREAM SMC-RNA Challenge, respectively (31). In a 2019 assessment of 23 fusion algorithms for cancer biology, both Arriba and STAR-Fusion ranked in the top three fastest and most accurate tools (32). *annoFuse* utilizes a four-step process (**Figure 1**) that is available with flexible functions to perform downstream functions such as merging, filtering, and prioritization of fusion calls from multiple fusion calling algorithms on single or batch samples.

**Figure 1.**
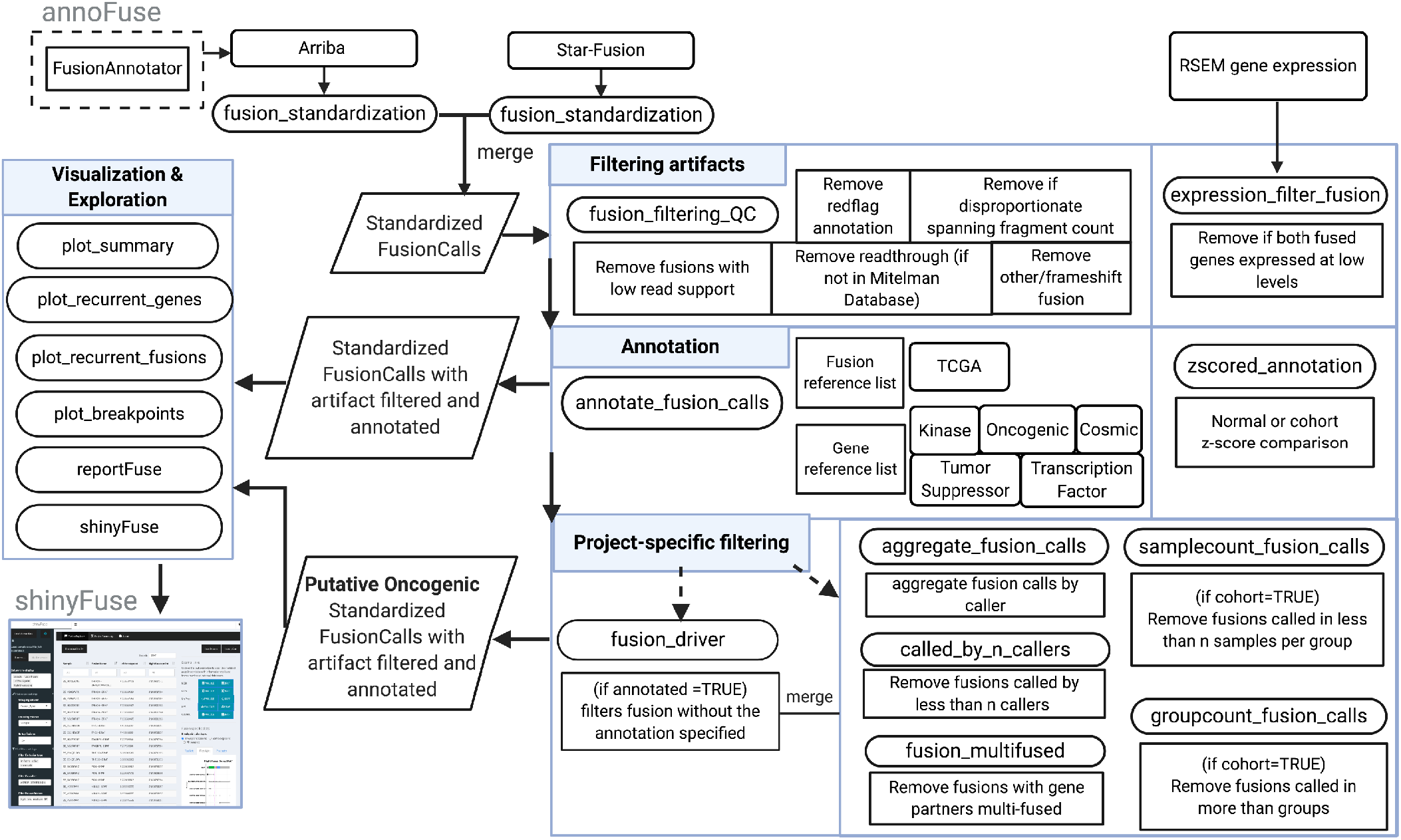
Graphical representation of the *annoFuse* pipeline. RNA-seq data processed through STAR-RSEM and fusion calls generated by Arriba v1.1.0 and/or STAR-Fusion 1.5.0 are inputs for the pipeline. The *fusion_standardization* function standardizes calls from fusion callers to retain information regarding fused genes, breakpoints, reading frame information, as well as annotation from FusionAnnotator. Standardized fusion calls use *fusion_filtering_QC* to remove false positives such as fusions with low read support, annotated as read-throughs, found in normal and gene homolog databases and remove non-expressed fusions using *expression_filter_fusion*. Calls are annotated with *annotate_fusion_calls* to include useful biological features of interest (eg. Kinase, Tumor suppressor etc.) Project-specific filtering captures recurrent fused genes using functions to filter (shown in boxes) as well as putative driver fusion. Outputs available from *annoFuse* include TSV files of annotated and prioritized fusions, a PDF summary of fusions, recurrently-fused gene/fusion plots, and an HTML report. Finally, users can explore fusion data interactively using shinyFuse. (Created with BioRender.com).

### RNA Expression and Fusion Calls

Currently, *annoFuse* is compatible with fusion calls generated from Arriba v1.1.0 (33) and/or STAR-Fusion 1.5.0 (14). Both tools utilize aligned BAM and chimeric SAM files from STAR as inputs and STAR-Fusion calls are annotated with GRCh38_v27_CTAT_lib_Feb092018.plug-n-play.tar.gz, which is provided in the STAR-fusion release. Arriba should be provided with strandedness information, or set to auto-detection for poly-A enriched libraries. Additionally, the blacklist file, blacklist_hg38_GRCh38_2018-11-04.tsv.gz contained in the Arriba release tarballs, should be used to remove recurrent fusion artifacts and transcripts present in healthy tissue. An expression matrix with FPKM or TPM values is also required; the matrix should have a column *“GeneSymbol* following the same gene naming convention as found in fusion calls.

### Fusion Call Preprocessing

We leveraged the fact that STAR-Fusion uses FusionAnnotator as its final step and thus, require all fusion calls be annotated with FusionAnnotator v. 0.2.0 to contain the additional column, *“annots”*. Fusion calls for all samples from both algorithms should be merged into a single TSV. Users will also provide a sample ID column (named “Sample” or other character vector specified by the user) for each merged file, which will be used to denote fusion calls per sample as well as identify recurrent fusions in cohorts of samples.

### annoFuse Steps

#### Step 1: Fusion Standardization

To obtain a standardized format for fusion calls from multiple fusion calls we use the *fusion_standardization* function to format caller specific output files to a standardizedFusionCalls format defined in the package README. *fusion_standardization* allows users to standardize fusion calls from multiple callers.

#### Step 2: Fusion Filtering

Events such as polymerase read-throughs, mis-mapping due to gene homology, and fusions occurring in healthy normal tissue confound detection for true recurrent fusion calls and false positives for genes considered as oncogenic, tumor suppressor or kinases in some cases. In this step, we filter the standardized fusion calls to remove artifacts and false positives (**Table 2**) using the function *fusion_filtering_QC*. The parameters are flexible to allow users to annotate and filter the fusions with *a priori* knowledge of their call set. For example, since the calls are preannotated with FusionAnnotator, the user can remove fusions known to be red-flags as annotated with any of the following databases GTEx_recurrent_STARF2019, HGNC_GENEFAM, DGD_PARALOGS, Greger_Normal, Babiceanu_Normal, BodyMap, and ConjoinG. This is done using the parameter, *artifact_filter = “GTEx_recurrent_STARF2019 | DGD_PARALOGS | Normal | BodyMap”*. Of note, we decided not to remove genes annotated in HGNC_GENEFAM, as this database contains multiple oncogenes and their removal resulted in missed true fusions using our validation truth set. Likewise, we retained fusions annotated with ConjoinG, as these may represent true chimeric RNA and protein products from adjacent genes, but are a separate class from read-through events (34). Read-throughs annotated by any algorithm can also be removed at this step by using the parameter *“readthroughFilter=TRUE”*. During validation, we observed the real oncogenic fusion, *P2RY8-CRLF2 (35,36)*, annotated as a read-through in acute lymphoblastic leukemia samples, therefore, we implemented a condition such that if a fusion is annotated as a read-through, but is present in the Mitelman cancer fusion database, we recover these fusions as true positive calls.

This function also allows users to flexibly filter out fusions predicted to be artifactual while retaining high-quality fusion calls using junction read support of ≥ 1 (default) and spanning fragment support of < 100 (default) reads compared to the junction read count, as disproportionate spanning fragment support indicates false positive calls (33). Finally, if both genes of the fusion are deemed not expressed < 1 FPKM or TPM (default), the fusion transcript calls can be removed using function *expressionFilterFusion*.

#### Step 3: Fusion Annotation

The *annotate_fusion_calls* function annotates standardized fusion calls and performs customizable fusion annotation based on user gene lists as input. If checkReciprocal == TRUE reciprocal status of fusions per sample_id is also provided. As a default setting, we provide lists of, and annotate gene partners as, oncogenes, tumor suppressor genes, and oncogenic fusions (**Table S1**).

The optional *zscored_annotation* function provides z-scored expression values from a user-supplied matrix such as GTEx or within cohort to compare samples with and without the fusion to look for over or under expression of fused genes compared to normal using a zscoreFilter. A cutoff of 2 (default) is set to annotate any score > 2 standard deviations away from the median as differentially-expressed. Researchers can then use this information to decide whether to perform additional downstream filtering.

### Single sample run

For single samples, we developed the *annoFuse_single_sample* function which performs fusion standardization of Arriba and STAR-Fusion calls, fusion filtering, and fusion annotation with user-provided gene and fusion reference lists.

### Project-Specific Filtering

Each study often requires additional downstream analyses to be performed once high-quality annotated fusion calls are obtained. We developed functions to enable analyses at a cohort (or project-level) and/or group-level (eg: histologies) designed to remove cohort-specific artifactual calls while retaining high-confidence fusion calls. The function *called_by_n_callers* annotates the number of algorithms that detected each fusion. We retained fusions with genes not annotated with the gene lists above (eg: oncogene, etc) that were detected by both algorithms as inframe or frameshift but not annotated as LOCAL_INVERSION or LOCAL_REARRANGEMENT by FusionAnnotator, as these could represent novel fusions. Additionally, *samplecount_fusion_call* identifies fusions recurrently called in (default ≥ 2) samples within each group. At the group-level, we add *groupcount_fusion_calls* (default ≥ 1) to remove fusions that are present in more than one type of cancer. At the sample level, *fusion_multifused* detects fusions in which one gene partner is detected with multiple partners (default ≥ 5), and we remove these as potential false positives. This enables *annoFuse* to scavenge back potential oncogenic fusions which may have otherwise been filtered. Separately, the function *fusion_driver* retains only fusions in which a gene partner was annotated as a tumor suppressor gene, oncogene, kinase, transcription factor, and/or the fusion was previously found in TCGA. Domain retention status for Gene1A (5’ gene) and Gene1B (3’ gene) for the given pfamIDs is also annotated and by default, we assess kinase domain retention status fusion-directed therapy often targets kinases. To further reduce the false positives and fusions containing pseudogenes from the cohort, we next filtered fusions using a cutoff of present in > 4 broad histologies after reviewing the fusion distributions within the OpenPBTA Project (**Additional File 1 : Figure S1**). Finally, potential driver fusions and scavenged back recurrent fusion sets are merged into a final set of putative oncogenic fusions.

### Fusion Domain Annotation

The *get_Pfam_domain* function in annoFuse provides domain annotation for each fused gene in standardized fusion calls. We used the UCSC pfamID Description database and domain location database (**Table S1**), along with bioMart (37,38) gene coordinates to get genomic locations of each domain in a gene. By identifying the breakpoint within the gene coordinate, we annotate each domain as described in (**Table S2**). This annotation provides domain retention information which enables prioritization to generate new hypotheses, validate fusion transcript functional impact, and or identify targeted therapeutic options.

### Visualization

Quick visualization of filtered and annotated fusion calls can provide information useful for review and downstream analysis. We provide the function *plot_summary*, which provides distribution of intra-chromosomal and inter-chromosomal fusions, number of in-frame and frameshift calls per algorithm, and distribution of gene biotypes, kinase group, and oncogenic annotation. If project-specific filtering is utilized, barplots displaying recurrent fusion and recurrently-fused genes can be generated using *plot_recurrent_fusions* and *plot_recurrent_genes*, respectively. *Finally, plot_breakpoints* can be used to generate all transcripts and breakpoints per gene to visualize the exon and domain retention resulting from the fusion (**Figure 4**).

### Interactive Fusion Exploration using shinyFuse

Depending on the size of the dataset, the prioritized fusions from *annoFuse* may still contain a considerable amount of information in need of further processing to efficiently extract biological insights. To facilitate this, we developed a web-based application, *shinyFuse*, in the R/Shiny framework, to assist users in performing drill-down operations while interacting with their fusion results. This feature is included in both the *annoFuse* package as well as a standalone server (http://shiny.imbei.uni-mainz.de:3838/shinyFuse/) to explore the results table, *PutativeDriverAnnoFuse.tsv*. Within the web interface, users can easily upload fusion calls from *annoFuse* or format a file from any fusion algorithm that generates minimal breakpoint location information. Prior to upload, users can choose to append additional columns containing sample descriptors and group information to take advantage of the interactive recurrence analysis feature. There are two major features of *shinyFuse: FusionExplorer* and *FusionSummary*.

### FusionExplorer

*FusionExplorer* allows users to interactively search, filter, visualize, and export the output of *annoFuse* through analysis of *PutativeDriverAnnoFuse.tsv*. Users can select specific fusion calls according to flexible combinations of filters (e.g. fusion type, caller count, spanning/junction fragments, spanning delta, confidence, and caller). Additionally, selecting a single fusion event (row) in the *FusionExplorer* tab generates breakpoint, exon, and protein domain plots tailored to transcripts for a specific fusion, sample, or for all samples. Full or filtered data, as well as plots, can be quickly downloaded from the application.

### FusionSummary

If a user has more than one cohort of samples and opts to utilize the project-specific filtering (eg: multiple samples from different types of cancer) or has multiple samples with factors by which they would like to group the data (eg: molecular subtype), these data can be explored using *FusionSummary*. The entire data table can be used for plotting, or it can first be filtered. The user can select a grouping column (factor), a counting column (usually patient-level), and the number of recurrent fusions to display in the plots. Recurrent fusions and recurrently-fused genes will be plotted and figures can be easily exported.

### Reproducible fusion analysis with reportFuse

Making results accessible and easier to interpret can play an essential role in reducing time and iterations required to extract actionable knowledge of large datasets, empowering a wide spectrum of collaboration partners. We acknowledge the importance of computational reproducibility (39,40) when generating analyses, and thus have created *reportFuse* with functionality to compile a full HTML report (using R Markdown). *reportFuse* is implemented using a template analysis containing multiple summary functions side by side with the code used to generate them. The report can be a valuable means for persistent storage and sharing of results with colleagues.

## Results and Discussion

### Technical validation of annoFuse

To assess our filtering strategy, we analyzed a subset of samples from TCGA and compared fusions retained with *annoFuse* filtering and prioritization to those deemed the final call set in a previously published analysis by The Fusion Analysis Working Group (13). A group of 160 samples were randomly selected BLCA (N = 10), BRCA (N = 11), CESC (N = 5), COAD (N = 11), ESCA (N = 5), GBM (N = 7), HNSC (N = 10), KIRP (N = 9), LGG (N = 9), LIHC (N = 9), LUAD (N = 5), LUSC (N = 11), OV (N = 9), PAAD (N = 8), PCPG (N = 2), PRAD (N = 14), SARC (N = 6), SKCM (N = 9), TGCT (N = 6), THCA (N = 4). We first ran STAR-Fusion, Arriba, and RSEM to generate fusion calls and gene expression values as described in OpenPBTA (16). Next, we standardized STAR-Fusion and Arriba fusion calls using *fusion_standardization*, then performed artifact and QC filtering using *fusion_filtering_QC*. Using a default spanningDelta (spanningFragCount - JunctionReadCount) of 10, *annoFuse* retained only 70% of the fusions in the final call set (**Table 3)**. Therefore, we visualized the distribution of spanningDelta (spanningFragCount - JunctionReadCount) across fusions called from the TCGA and PBTA cohorts to assess a cutoff for spanningDelta (**Additional file 1 : Figure S2**). We found that sensitivity reaches 96% at cutoff of 100 (**Additional file 1 : Figure S3** and **Table 3**) for TCGA 50 to 76 bp read length RNAseq data. Therefore, we have implemented a default spanningDelta of 100 and made this a customizable input parameter.

Few gene fusion “truth” sets exist and those that do consist of simulated data or synthetic fusions spiked into breast cancer cell lines or total RNA (31,32,41). We therefore utilized a recent study in which high-confidence fusions were reported in 244 patient-derived xenograft models from the Pediatric Preclinical Testing Consortium (PPTC) (42). A set of 27 fusions were molecularly validated from acute lymphoblastic leukemia (ALL) models in the PPTC dataset and we deemed this our “truth” set. We first ran Arriba on the PPTC dataset and determined that 23 of the 27 truth fusions were detected using only STAR-Fusion and/or Arriba (). Next, we used *annoFuse* to filter and prioritize putative oncogenic fusions. **Table 4** shows the performance of *annoFuse*, which retained all 23 true positive ALL fusions (100%). Interestingly, only 114 of 166 previously defined as high-confidence (putative oncogenic fusions) in (42) fusions were detected using STAR-Fusion and Arriba (23/27 within the “truth” set), implying gold standard algorithms alone still fail to capture the full landscape of gene fusions, reflecting that additional algorithms should be integrated into our workflow. Of the 114 fusions we detected, 110 (96%) were retained as putative oncogenic fusions using *annoFuse*. The four fusions *annoFuse* did not retain were removed with default the “read-through” filter, which can be turned off as an option.

### Case study with annoFuse, shinyFuse, and reportFuse using OpenPBTA

As proof of concept, we utilized RNA expression generated by STAR-RSEM (43) and fusion calls generated by Arriba v1.1.0 (33) and/or STAR-Fusion 1.5.0 (14) which were released as part of the Pediatric Brain Tumor Atlas (44). Briefly, RNA from fresh-frozen tissue was extracted and libraries were prepped and sequenced at 2×100 or 2×150 bp to an average of 200M+ total reads, and at least 60% of reads were required to map to the human genome for fusion analysis to proceed. Additional details can be accessed in the OpenPBTA manuscript (16). The algorithms were run as described in ***RNA Expression and Fusion Calls*.** The RNA expression and fusion workflows are publicly available within the Gabriella Miller KidsFirst GitHub repository (45).

Following fusion standardization, annotation, and filtration, we applied project-specific filtering to the OpenPBTA RNA-Seq cohort (n = 1,028 biospecimens from n = 943 patients). **Figure 2** is a sample summary PDF designed to give the user an overall glance of the fusion annotations and fusion characteristics within the cohort. From the OpenPBTA cohort, it is clear that there were predominantly more intra-chromosomal fusions called than inter-chromosomal fusions, even after filtering for read-through events (**Figure 2A**). While a low-grade astrocytic tumors are the major pediatric brain tumor subtype known to be enriched for gene fusions, it was surprising to observe a large number of fusions in diffuse astrocytic and oligodendroglial tumors and the project-specific utility of *annoFuse* allows researchers to further prioritize fusions. Histologies within the OpenPBTA project were classified according to broad WHO 2016 subtypes (46).

**Figure 2.**
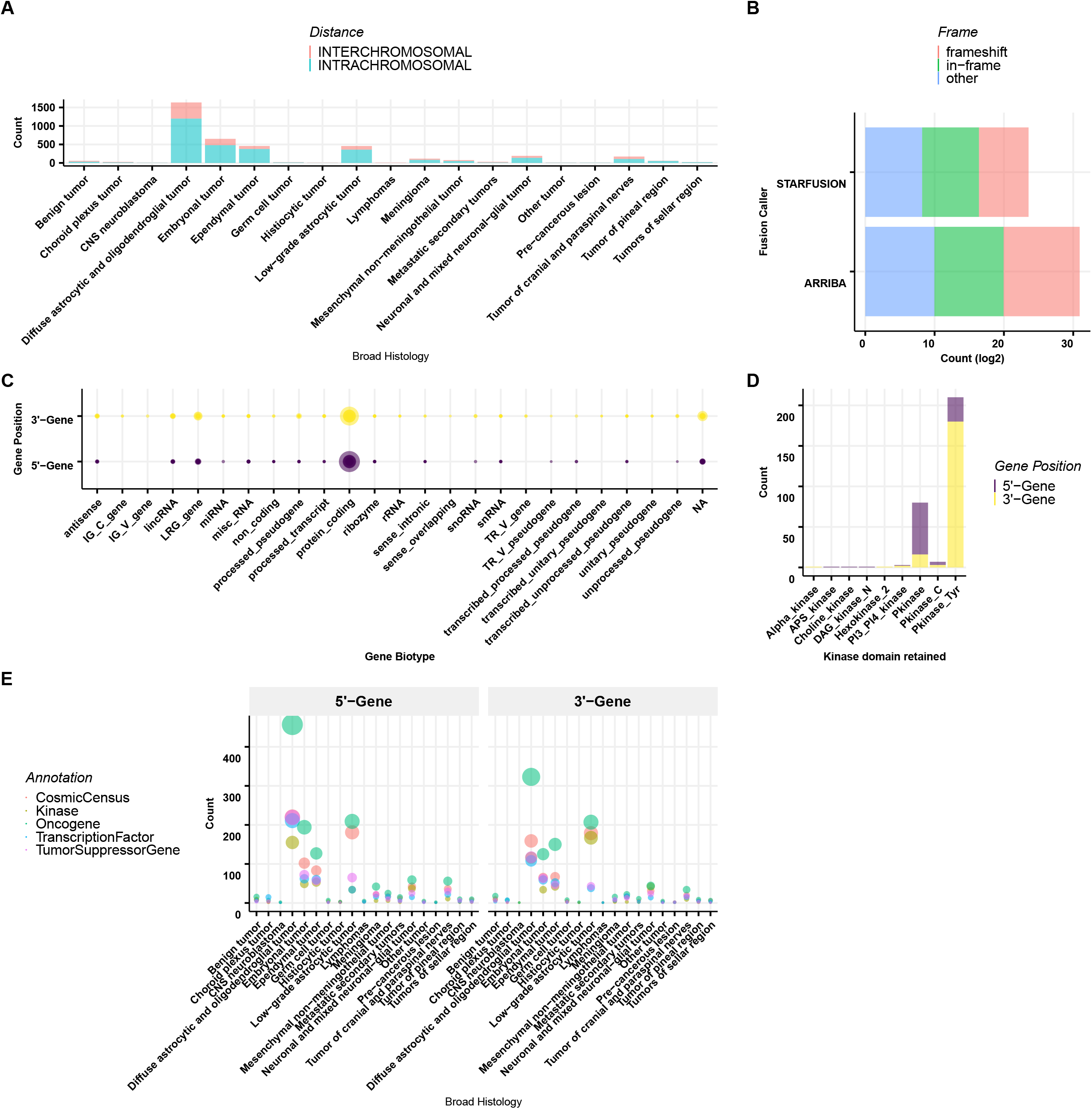
Fusion annotations generated by annoFuse. (A) Distribution of intra- and inter-chromosomal fusions across histologies. (B) Transcript frame distribution of fusions detected by Arriba and STAR-Fusion algorithms. (C) Bubble plot of gene partner distribution with respect to ENSEMBL biotype annotation (Size of circle proportional to number of genes). (D) Barplots representing the distribution of kinase groups represented in the PBTA cohort annotated by gene partner. (Alpha_kinase=Alpha-kinase family, Hexokinase_2=Hexokinase, PI3_PI4_kinase=Phosphatidylinositol 3- and 4-kinase, Pkinase=Protein kinase domain, Pkinase_C=Protein kinase C terminal domain,Pkinase_Tyr=Protein tyrosine kinase) (E) Bubble plot representing the distribution of fused genes as oncogenes, tumor suppressor genes, kinases, COSMIC, predicted and curated transcription factors (Size of circle proportional to number of genes). Genes belonging to more than one category are represented in each. In all panels except for B, fusion calls were merged from both STAR-Fusion and Arriba.

The number of in-frame and frameshift fusions per algorithm were roughly equivalent within each STAR-Fusion and Arriba fusion calls (**Figure 2B**). **Figure 2C** depicts the density of genes categorized by gene biotype (biological type), and as expected, the filtered and annotated calls were enriched for biologically-functional fusions; the majority of gene partners are classified as protein-coding. The majority of gene partners were annotated as tyrosine kinase (TK) or tyrosine kinase-like (TKL) (**Figure 2D**). In **Figure 2E**, the user can explore the biological and oncogenic relevance of the fusions across histologies. Of the fusions harboring kinase domains, we found that the majority of 3’ partners retained kinase domains, supporting these fusions as functionally relevant (**Additional file 1 : Figure S4**).

Following project-specific filtering, we observed *KIAA1549-BRAF* fusions as the most recurrent in-frame fusion in our cohort (n = 109/898), since *KIAA1549-BRAF* expressing low-grade astrocytic tumors comprise the largest representative histology in the OpenPBTA cohort (n = 236/898). *C11orf95-RELA* was predominant in ependymal tumors (n = 25/80), as expected in supratentorial ependymomas (47). Other expected recurrent oncogenic fusions obtained through *annoFuse* were *EWSR1-FLI1* in CNS Ewing sarcomas (48), and *KANK1-NTRK2, MYB-QKI*, and *FAM131B-BRAF* in low-grade astrocytic tumors (4,49) (**Figure 3A**). In addition to recurrent fusions, we also detect recurrently-fused genes to account for partner promiscuity. This enables us to see a broader picture of gene fusions, specifically within diffuse astrocytic and oligodendroglial tumors, in which we see fusions prevalent in *ST7, MET, FYN, REV3L, AUTS2*, and *ROS1*, and meningiomas, in which *NF2* fusions are common. (**Figure 3B**). Next, we added functionality to visualize domain information (**Figure 4**) to quickly scan for domains retained and lost across the dataset. The putative oncogenic fusion table from this analysis is both available in the vignette and on the standalone Shiny server as a demo dataset for *shinyFuse*, enabling both free exploration of the data as well as reproducible generation of Figures 3 and 4. Finally, we use *reportFuse* to include a reproducible HTML report which generates all output tables and figures for this manuscript.

**Figure 3.**
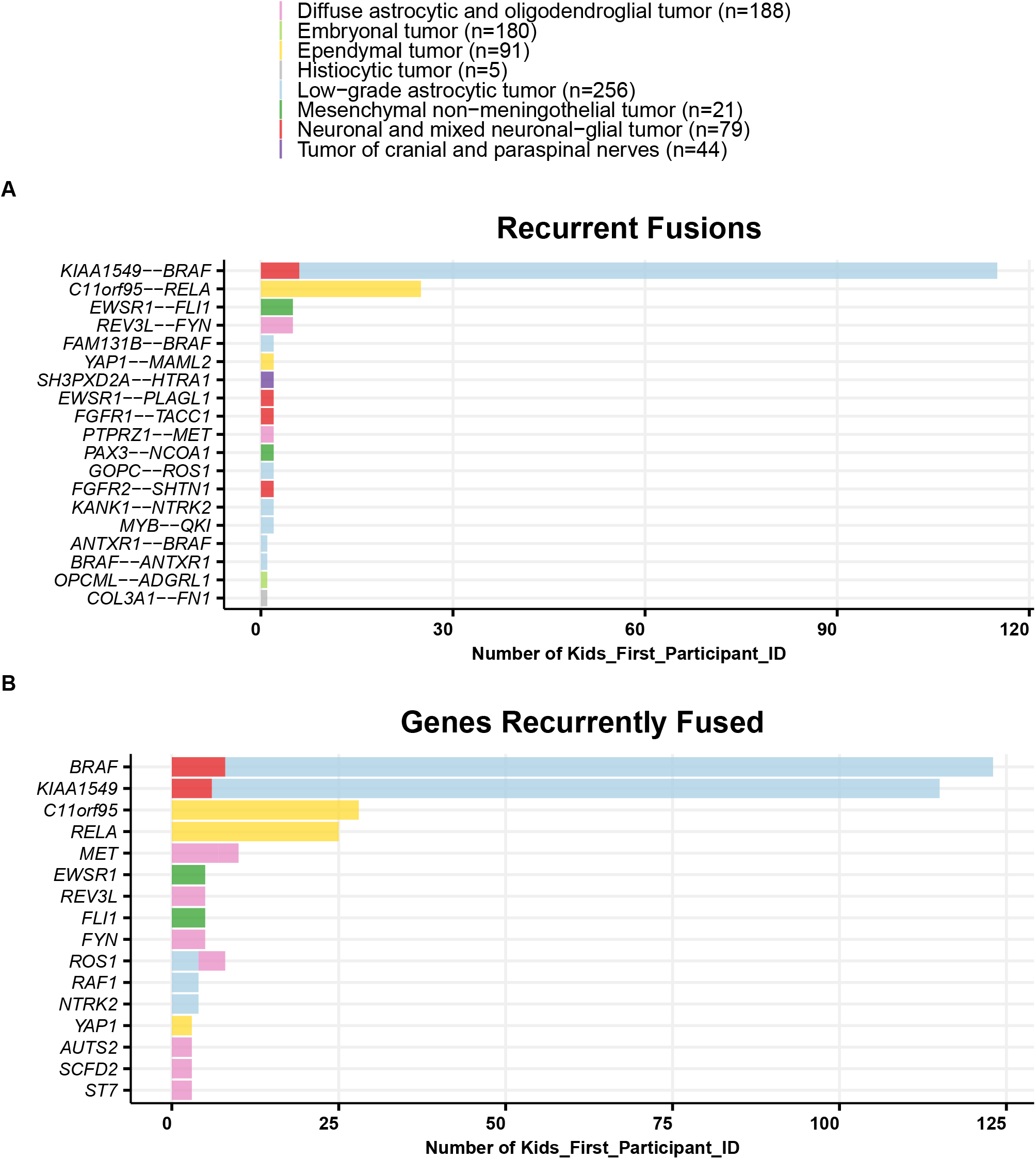
Recurrent fusion plots generated by annoFuse. Bar plots as representative of histology showing recurrent fusion calls by number of patients (A) and recurrently-fused genes by number of patients (B) after filtering and annotation.

**Figure 4.**
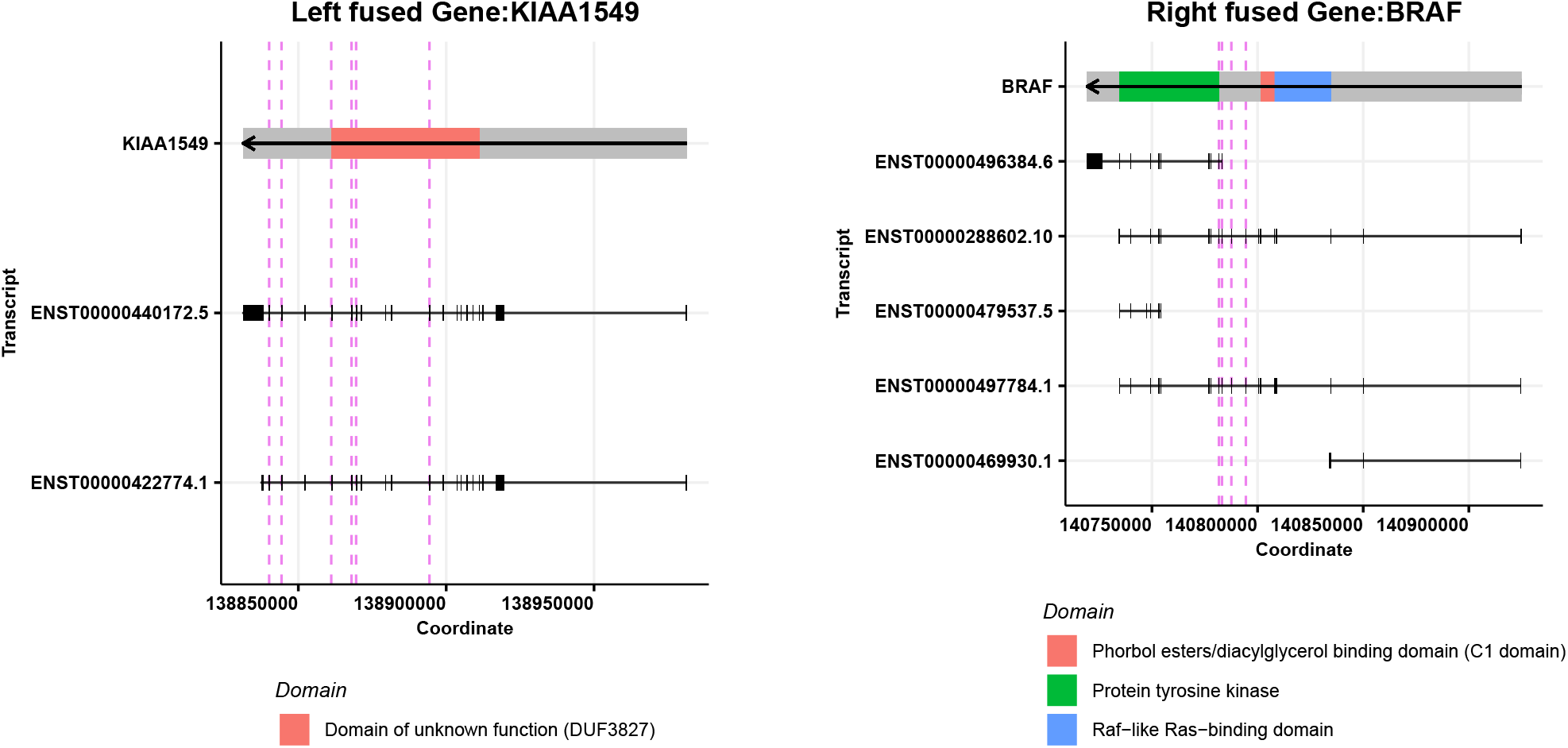
Breakpoint distribution for *KIAA1549-BRAF* fusion. Displayed are each fusion gene, all known transcripts, and their genomic coordinates. Red dotted lines in each gene panel are the locations of breakpoints detected for *KIAA1549* (3’) and *BRAF* (5’) compiled from both Arriba and STAR-Fusion. Strand information is depicted with an arrow in the gene and domain as colored boxes. Black boxes represent exons for each transcript.

The few openly-available fusion annotation and/or prioritization tools summarized in **Table 1** perform only a subset of the functionalities provided in *annoFuse* and *shinyFuse*. In addition, the majority are no longer maintained, nor are directly compatible with Arriba, the tool that won the SMC-RNA DREAM challenge for speed and accuracy in 2019. For example, Oncofuse (50), Pegasus (51), chimera (52), and co-Fuse (53) have not been updated in two or more years, and as a result, these tools lack compatibility with newer and improved fusion algorithms. The chimeraviz R package (54) is well-maintained and compatible with nine fusion algorithms, but only performs visualizations of fusions, thus prioritization using database annotation, readthrough and expression is not possible using this tool. Four tools are algorithm agnostic, yet perform only specific aspects of annotation and prioritization. Oncofuse (50) and Pegasus (51) analyze oncogenicity of fusions, and chimera (52) and FusionHub (55) require Oncofuse (50) to be run as a first step. FusionHub (55) is a web-based tool which enables annotation of fusions with 28 databases, however, is not programmatically scalable. FusionAnnotator (56) determines the presence of fusions in 15 cancer-associated databases, oncogene lists, and seven databases for fusions not relevant in cancer, but does not perform prioritization. ConFuse performs filtering and prioritization, but is not algorithm-agnostic; it only works on fusion calls from deFuse (57). AGFusion (58) annotates protein domains, and Fusion Pathway (59) utilizes fusion and protein domain annotations in gene set enrichment analysis (GSEA) to infer oncogenic pathway association. When used exclusively, none of these tools flexibly perform fusion annotation and prioritization. Furthermore, none enable the interactive exploration and visualization capabilities that we provide with *shinyFuse*. Instead, we leveraged the algorithm agnostic capabilities of FusionAnnotator to pre-annotate fusion input from STAR-Fusion and Arriba to create *annoFuse*, an all-in-one tool that performs fusion annotation, oncogenic prioritization, recurrence analysis, visualization, and exploratory fusion analysis.

By integrating FusionAnnotator with functionality of the current gold standard algorithms STAR-Fusion and Arriba, we were able to improve the aforementioned tools’ capabilities by meeting the current demands of the research community. We provide the user with flexible filtering parameters and envision *annoFuse* will be used to quickly filter sequencing artifacts and false positives, as well as further annotate fusions for additional biological functionality (eg: kinases, transcription factors, oncogenes, tumor suppressor genes) to prioritize fusion calls in a cancer cohort. Additionally, users can opt to simply annotate and filter artifacts or use *annoFuse* to functionally prioritize fusions as putative oncogenic drivers. During the prioritization steps, we filter based on genes with cancer relevance (see biological functionality list above) and perform analysis of fusion and fused-gene recurrence to create a stringently filtered, prioritized list of fusions likely to have oncogenic potential.

### Limitations and Future Directions

It is worth noting that *annoFuse* cannot correct fusion calls derived from low-quality RNA-Seq data, thus onus is on the user to design the extraction, library preparation, and sequencing portions of the experiment to enable production of high-quality data. We prioritized Arriba and STAR-Fusion calls as input to *annoFuse* due to their performance in the SMC-RNA DREAM challenge and we recommend utilizing both algorithms prior to analysis with *annoFuse*. In addition, users can provide their own informed cutoffs for read support and annotation filters to enable appropriate prioritization of oncogenic fusions. We are actively adding compatibility with additional fusion algorithms currently used by the community, such as deFuse, FusionCatcher, SOAPfuse, and Jaffa to further increase the applicability of *annoFuse*. As an additional feature, we plan to add expression-based comparison of genes between fused samples, normal, and within a histology or cohort. Future features could link domain retention to drug databases to predict fusion-directed targeting strategies.

## Conclusions

Gene fusions provide a unique mutational context in cancer in which two functionally-distinct genes could be combined to function as a new biological entity. Despite showing great promise as diagnostic, prognostic, and therapeutic targets, translation in the oncology clinic is not yet accelerated for gene fusions. This has been partly due to limited translation of the large number of computationally-derived fusion results into biologically meaningful information. In our efforts to address this, we introduce *annoFuse*, an R Package to annotate and prioritize putative oncogenic RNA fusions and *shinyFuse*, an algorithm-agnostic web application for interactive fusion exploration and plotting. We include a cancer-specific workflow to find recurrent, oncogenic fusions from large cohorts containing multiple cancer histologies. The filtering and annotation steps within *annoFuse* enable users to integrate calls from multiple algorithms to improve high-confidence, consensus fusion calling. The lack of concordance among algorithms as well as variable accuracy with fusion truth sets (2,32) adds analytical complexity for researchers and clinicians aiming to prioritize research or therapies based on fusion findings. Through *annoFuse*, we add algorithm flexibility and integration to identify recurrent fusions and/or recurrently-fused genes as novel oncogenic drivers. Within the package, *shinyFuse* and *reportFuse* deliver interactive and reproducible analysis options to efficiently extract knowledge from the outputs of the *annoFuse* workflow. We expect *annoFuse* to be broadly applicable to cancer datasets and empower researchers and clinicians to better inform preclinical studies targeting novel, putative oncogenic fusions and ultimately, aid in the rational design of therapeutic modulators of gene fusions in cancer.

## Availability and requirements

**Project name:** *annoFuse:* an R Package to annotate, prioritize, and interactively explore putative oncogenic RNA fusions

**Project home page**: https://github.com/d3b-center/annoFuse

**Project web application (shinyFuse): http://shiny.imbei.uni-mainz.de:3838/shinyFuse/**

**Operating system(s:** Platform independent

**Programming language:** R (>=4.0.0)

**License:** MIT

**Any restrictions to use by non-academics:** None

## List of abbreviations

ALL: Acute Lymphoblastic Leukemia
BAM: Binary Alignment Map
COSMIC: Catalogue Of Somatic Mutations In Cancer
CNS: Central Nervous System
DGD_PARALOGS: Duplicated Genes Database annotated paralogs
GSEA: Gene Set Enrichment Analysis
HGNC_GENEFAM: HGNC annotated gene family
FPKM: Fragments Per Kilobase Million
OpenPBTA: Open Pediatric Brain Tumor Atlas
PI3_PI4_kinase: Phosphatidylinositol 3- and 4-kinase
Pkinase: Protein kinase domain
Pkinase_C: Protein kinase C terminal domain
Pkinase_Tyr: Protein tyrosine kinase
PPTC: Pediatric Preclinical Testing Consortium
RNA: Ribonucleic Acid
SAM: Sequence Alignment Map
SMC-RNA: Somatic Mutation Calling RNA DREAM Challenge (SMC-RNA)
TCGA: The Cancer Genome Atlas
TSV: Tab Separated Value
TPM: Transcripts Per Kilobase Per Million
WHO: World Health Organization

## Declarations

### Ethics approval and consent to participate

Not applicable.

### Consent for publication

Not applicable.

### Availability of data and materials

All brain tumor raw data are available by download from the Gabriella Miller Kids First Data Resource Center with a data access agreement through the Children’s Brain Tumor Network and processed data (release-v16-20200320) available by download through the OpenPBTA project’s GitHub repository: https://github.com/AlexsLemonade/OpenPBTA-analysis. PPTC (Accession Number phs001437.v1.p1) and TCGA (Accession Number phs000178.v1.p1) RNA-Sequencing data are available by download from dbGAP with a data access agreement.

### Authors’ contributions

Conceptualization: KSG, JMM, KSR, PR, JLR, JNT, FM

Methodology: KSG, KSR, PR, JLR, JNT, MAB, BZ, YZ, NAC Software: KSG, JLR, JNT, FM

Validation: KSG, JLR, JNT, FM

Formal Analysis: KSG, JNT

Investigation: KSG, JLR, FM

Resources: JLR, PBS, ACR, KS

Data Curation: KSG, YZ, JLR, MAB, BZ

Writing - Original Draft: KSG, JLR

Writing - Review & Editing: KSG, JLR, PJ, ASN, JNT, FM

Visualization: KSG, JLR, FM

Supervision: JLR, JNT

Funding Acquisition: JLR, ACR, PBS, JMM, JNT, FM

## Acknowledgements

We would like to thank Brian J. Haas (Broad Institute), Alexander Dobin (Cold Spring Harbor Laboratory) and Sebastian Uhrig (German Cancer Research Center) for helpful discussions throughout the development of *annoFuse*. We acknowledge the Children’s Brain Tumor Network and the Pacific Pediatric Neuro-Oncology Consortium for generating RNA-Seq data from pediatric brain tumor tissue and the Gabriella Miller Kids First Data Resource Center for processing and hosting the data. Finally, we graciously thank the patients and families for donating tumor tissue, without which, this research would not be possible.

## Additional file 1

**Figure S1. Fusions found in more than 1 histology.** Barplots represent the number of histologies in which each fusion was observed. Dotted line represents the cut off (> 4 counts) used to remove potential false positives and fusions containing pseudogenes.

**Figure S2. Distribution of spanningDelta for *annoFuse* prioritized fusions from TCGA and PBTA cohorts**

The spanningFragCountFilter and mean are plotted for each filtering cutoff of 10, 20, 30, 40, 50, 100, 150, and 200 spanningDelta for TCGA (N = 160 samples) (A) and PBTA (N = 1028 samples) (B) fusions.

**Figure S3 : Sensitivity of TCGA fusions retained by annoFuse.**

At a spanningDelta of 100, *annoFuse* achieved a sensitivity of 96.35% for fusions in the TCGA final call set. The x-axis represents the cutoff for spanningFragCountFilter used and y-axis represents the sensitivity of the fusions retained after *fusion_filtering_QC*.

**Figure S4. Distribution of kinase genes fused in 5’ and 3’ genes per histologies.** For each broad histology, pie charts represent the percentage of fusions in which Gene1A (5’) or Gene1B (3’) retain their kinase domains.

**Table 1. Available fusion annotation and prioritization tools.** List of ten openly-available fusion annotation and prioritization software tools, compared to *annoFuse*. Only AGFusion, FusionAnnotator, Fusion Pathway, and certain functions of FusionHub are algorithm agnostic, and most algorithms require outdated fusion algorithm input.

**Table 2. Fusion filtering and annotation criteria.** Fusion filtering criteria were developed to gather high quality recurrent fusion calls while retaining fusions containing oncogenes and/or tumor suppressor genes. Filtering is divided into 3 types 1) QC: filters known causes of false positives. 2) Gene-list: retains additional fusions in genes and fusions of interest list. 3) Recurrence: filters out non-recurrent fusions in genes not annotated as putative oncogenic. Annotation lists are also described.

**Table 3. Sensitivity of TCGA fusion calls.** Fusion standardization and fusion artifact filtering was conducted on a subset of TCGA samples and compared to published filtered fusion calls from The Fusion Analysis Working Group. SpannigFragCountFilter cutoffs of 10, 20, 30, 40, 50, 100, 150, and 200 were assessed to determine sensitivity of *annoFuse* prioritized fusion calls.

**Table 4. Validation of *annoFuse* prioritization using PPTC PDX fusion calls.** Retention of high-confidence, putative oncogenic calls averaged 96% across the entire PPTC PDX dataset and was 100% for the ALL truth set (ALL = acute lymphoblastic leukemia). Column 1 = PPTC histology, Column 2 = fusion calls from STAR-Fusion, FusionCatcher, deFuse, and SOAPFuse which were filtered and reported as high-confidence in the PPTC dataset, Column 3 = PPTC reported fusions detected from STAR-Fusion and Arriba, Column 4 = Fusions retained following *annoFuse* filtering, Column 5 = Percent of fusions retained after applying *annoFuse*.

**Table S1. Sources for annotation of fusions and gene partners.** Listed are annotations utilized by *annoFuse* for prioritization of oncogenic gene fusions, along with reference files and their sources.

**Table S2. Domain retention conditions for annotation.** Listed are conditions for domain retention status for Gene1A (5’ gene) and Gene1B (3’ gene). Domain retention status for Gene1A depends on the direction of the left breakpoint with respect to the domain and the gene body and domain retention status for Gene1B depends on the direction of the right breakpoint with respect to domain and gene body.

