## Supplementary Figures for "annoFuse: an R Package to annotate, prioritize, and interactively explore putative oncogenic RNA fusions"

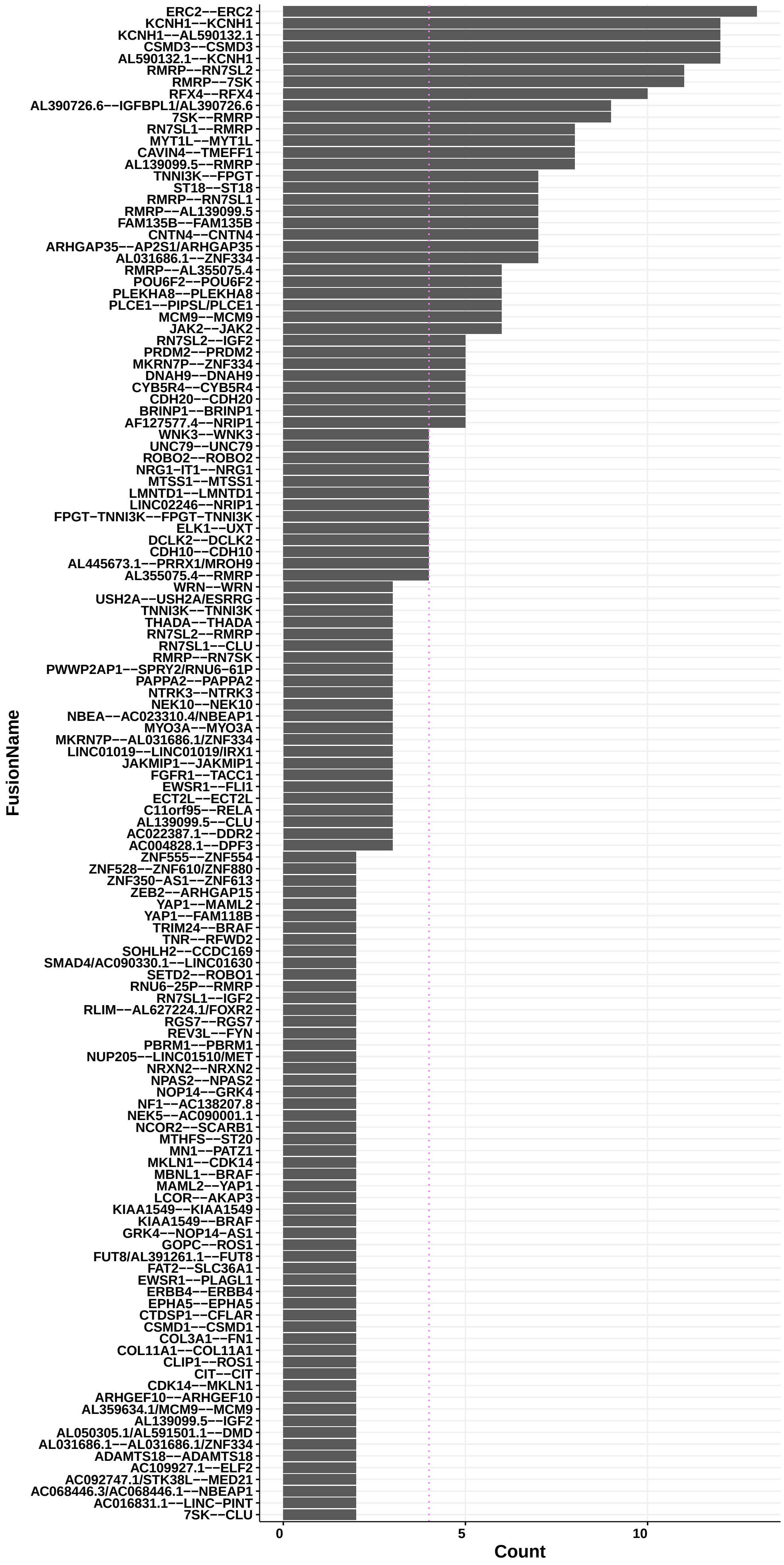


**Figure S1:** Fusions found in more than 1 histology

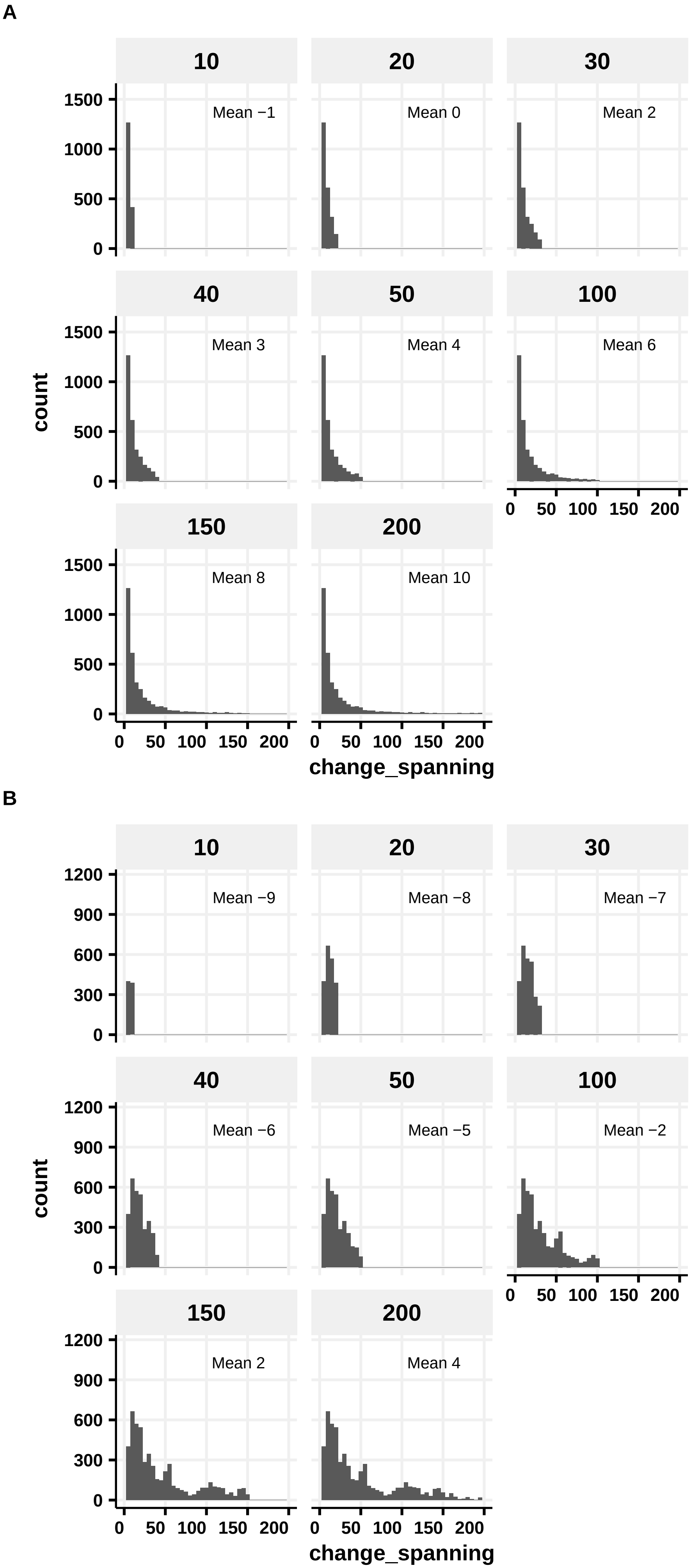


**Figure S2:** Distribution of spanningDelta for annoFuse prioritized fusions from TCGA and PBTA cohorts

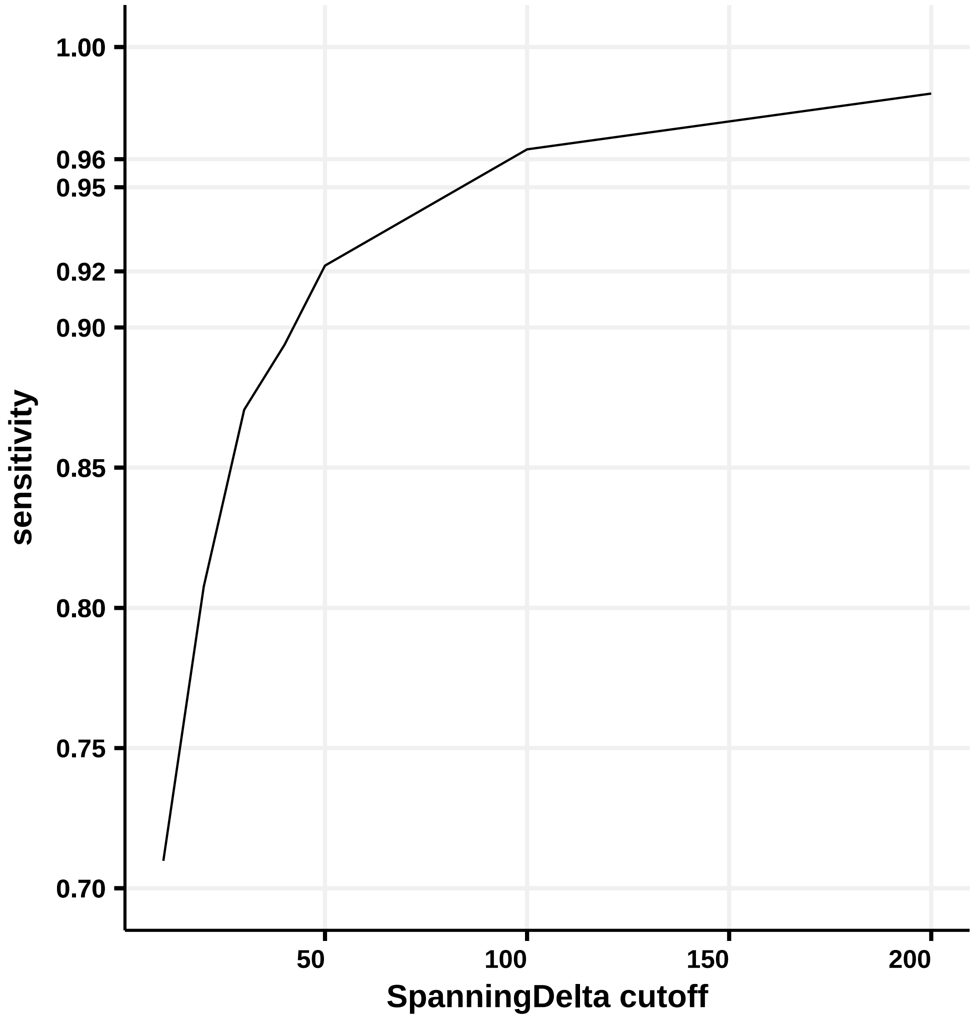


**Figure S3:** Sensitivity of TCGA fusions retained by annoFuse

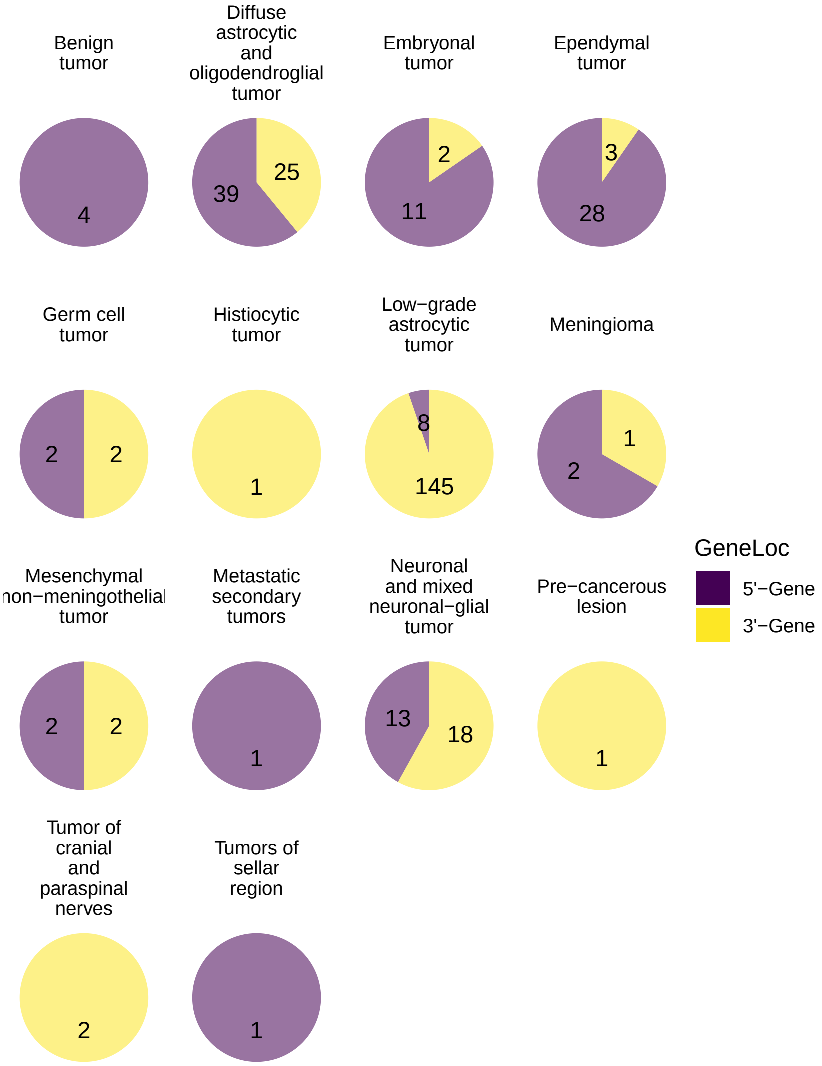


**Figure S4:** Distribution of kinase genes fused in 5' and 3' genes per histologies
